# Taking a Dive: Experiments in Deep Learning for Automatic Ontology-based Annotation of Scientific Literature

**DOI:** 10.1101/365874

**Authors:** Prashanti Manda, Lucas Beasley, Somya D. Mohanty

## Abstract

Text mining approaches for automated ontology-based curation of biological and biomedical literature have largely focused on syntactic and lexical analysis along with machine learning. Recent advances in deep learning have shown increased accuracy for textual data annotation. However, the application of deep learning for ontology-based curation is a relatively new area and prior work has focused on a limited set of models.

Here, we introduce a new deep learning model/architecture based on combining multiple Gated Recurrent Units (GRU) with a character+word based input. We use data from five ontologies in the CRAFT corpus as a Gold Standard to evaluate our model’s performance. We also compare our model to seven models from prior work. We use four metrics - Precision, Recall, F1 score, and a semantic similarity metric (Jaccard similarity) to compare our model’s output to the Gold Standard. Our model resulted in a 84% Precision, 84% Recall, 83% F1, and a 84% Jaccard similarity. Results show that our GRU-based model outperforms prior models across all five ontologies. We also observed that character+word inputs result in a higher performance across models as compared to word only inputs.

These findings indicate that deep learning algorithms are a promising avenue to be explored for automated ontology-based curation of data. This study also serves as a formal comparison and guideline for building and selecting deep learning models and architectures for ontology-based curation.

## II. Introduction

Ontology-based data representation has been widely adopted in data intensive fields such as biology and biomedicine due to the need for large scale computationally amenable data [1]. However, the majority of ontology-based data generation relies on manual literature curation - a slow and tedious process [2]. Natural language and text mining methods have been developed as the solution for scalable ontology-based data curation [3, 4].

One of the most important tasks for annotating scientific literature with ontology concepts is Named Entity Recognition (NER). In the context of ontology-based annotation, NER can be described as recognizing ontology concepts from text [5]. Outside the scope of ontology-based annotation, NER has been applied to biomedical and biological literature for recognizing genes, proteins, diseases, etc [5].

The large majority of ontology driven NER techniques rely on lexical and syntactic analysis of text in addition to machine learning for recognizing and tagging ontology concepts [3, 4, 6]. In recent years, deep learning has been introduced for NER of biological entities from literature [7, 8, 9, 10, 11]. However, the majority of prior work has focused on a limited set of models, particularly, the Long Short-Term Memory (LSTM) model (e.g. [7]).

Here, we present a new deep learning architecture that utilizes Gated Recurring Units (GRU) while taking advantage of word and character encodings from the annotation training data to recognize ontology concepts from text. We evaluate our model in comparison to 7 deep learning models used in prior work to show that our model outperforms the state of art at the task of ontology-based NER.

We use the Colorado Richly Annotated Full-Text (CRAFT) corpus [12] as a Gold Standard reference to develop and evaluate the deep learning models. The CRAFT corpus contains 67 open access, full length biomedical articles annotated with concepts from several ontologies (such as Gene Ontology, Protein Ontology, Sequence Ontology, etc.). We use four metrics - 1) Precision, 2) Recall, 3) F-1 Score and 4) Jaccard semantic similarity to compare each model’s performance to the Gold Standard.

Precision and Recall are traditionally used to assess the performance of information retrieval systems. However, these metrics do not take into account the notion of partial information retrieval which is important for ontology-based annotation retrieval. Sometimes, an NLP system might not retrieve the same ontology concept as the gold standard but a related concept (sub-class or super-class). To assess the performance of the NLP system accurately, we need semantic similarity metrics that can measure different degrees of semantic relatedness between ontology concepts [13]. Here, we use Jaccard similarity to compare annotations from each deep learning model to the gold standard. Jaccard similarity assesses similarity between two ontology terms based on the ontological distance between them - the closer two terms are, the more similar they are considered to be [13].

## III. Related Work

The application of deep learning for ontology-based Named Entity Recognition is a nascent area with relatively little prior work. Habibi et al. [9] studied entity recognition on biomedical literature using long short-term memory network-conditional random field (LSTM-CRF) and showed that the method outperformed other NER tools that do not use deep learning or use deep learning methods without word embeddings. Lyu et al. [10] also explored LSTM based models enhanced with word and character embeddings. They do not evaluate other deep learning models but present results only based on LSTM with word embeddings. Wang et al. [11] also propose a LSTM based method for recognizing biomedical entities from literature. Similar to the above studies, Wang et al. show that a bidirectional LSTM method used with Conditional Random Field (CRF) and word embeddings outperforms other methods. The striking difference between these prior studies and our work here is that the majority of prior literature focuses on LSTM based methods along with CRF and word embeddings. The potential of other deep learning models such as Recurrent Neural Networks, Gated Recurrent Units, etc., at the task of ontology-based NER remains unexplored presenting a unique need and opportunity. Our study aims to fill this knowledge gap. In addition, all the above studies focus on non-ontology based NER for entities such as genes, disease names, etc. In contrast, our study’s focus is on recognizing ontology concepts within text.

## IV. Methods

### A. Data Preprocessing

Annotation files for the 67 papers in CRAFT were cleaned to remove punctuation symbols (except for period at the end of sentences), special symbols, and non-ASCII characters. Annotations for GO, CHEBI, Cell, Protein, and Sequence ontologies were converted from the cleaned files to separate ontology-specific text files that represent the presence or absence of ontology terms. For each ontology, every sentence containing at least one annotation from that ontology was represented using two lines in the ontology-specific text file. The first of these two lines contained an array with each word in the sentence. The second contained an ordered encoding corresponding to words in the first line. These encodings could be an ontology concept ID if the corresponding word was annotated in CRAFT or an ‘*O*’ if the corresponding word was not annotated.

For example, the sentence “Rod and cone photoreceptors subserve vision under dim and bright light conditions respectively” where the word “vision” was annotated to GO ID “*GO:0007601 (perception of sight)*” would be represented using the two lines below:

- *[‘Rod’, ‘and’, ‘cone’, ‘photoreceptors’, ‘subserve’, ‘vision’, ‘under’, ‘dim’, ‘and’, ‘bright’, ‘light’, ‘conditions’, ‘respectively’]*
- *[‘O’, ‘O’, ‘O’, ‘O’, ‘O’, ‘GO:0007601’, ‘O’, ‘O’, ‘O’, ‘O’, ‘O’, ‘O’, ‘O’]*

Annotations to single words (unigrams) were only included in these preprocessed files. So, if an annotation was made in CRAFT to a phrase containing more than one word, it was ignored in the preprocessed data.

### B. Performance evaluation metrics

Precision, Recall, F1-score, and Jaccard similarity were used to evaluate the performance of the models. The Jaccard similarity (*J*) of two ontology concepts (in this case, annotations) (*A*, *B*) in an ontology is defined as the ratio of the number of classes in the intersection of their subsumers over the number of classes in their union of their subsumers [13].

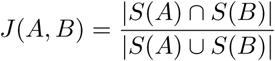

where *S*(*A*) is the set of classes that subsume A. Jaccard similarity ranges from 0 (no similarity) to 1 (exact match).

### C. Deep learning models

Below, we describe four deep learning models - Multilayer perceptrons, Recurrent Neural Networks, Long Short-Term Memory, and Gated Recurrent Units. Next, we describe three architectures - window based, word based, and character-word based that can be used in conjunction with the above models. Finally, we describe our new model that combines character-word based architecture with Gated Recurrent Units and six models used in prior work.

#### 1) Multi-Layer Perceptron (MLP)

A Multi-Layer Perceptron (MLP) [14] is a feed-forward deep-neural network model which consists of an input, single/multiple hidden, and an output layer, each consisting of a number of perceptrons. A single perceptron computes the output as 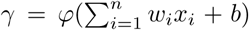, where *w* is the weight vector, *x* is the provided input, *b* is the bias, and *ϕ* is the activation function. The weights and biases of each perceptron in the layers are adjusted using back-propagation to minimize prediction error

#### 2) Recurrent Neural Network

A Recurrent Neural Network (RNN) [15] is an adaption of feed-forward neural networks, where history of the input sequence is taken into consideration for future prediction. Given an input sequence < *x*_0_*,x*_1_*,x*_2_, ⋯ *x_i_* >, the hidden state (*h_t_*) of an RNN is updated as follows:

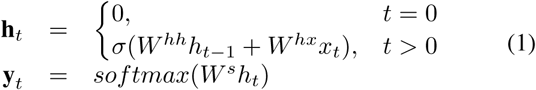

where, *x_t_* is the input provided to the hidden state *h_t_* at time *t* which is updated using a *sigmoid* function *σ*. *σ* is calculated over the previous time state of the network given by *h_t_*_−1_ and current input *x_t_*. *W_hh_*, *W_hx_*, and *W^s^* are the weights computed over training. The network can then produce an output prediction < *y*_0_*,y*_1_*,y*_2_, ⋯ *y_j_* > using a *softmax* function on the hidden state *h_t_*.

A bidirectional Recurrent Neural Network (BiRNN) is an RNN where the input data is fed to the neural network two times - once in forward and again in reverse order.

#### 3) Long-Short Term Memory

While RNNs are effective in learning temporal patterns, they suffer from a vanishing gradient problem where long term dependencies are lost. A solution to the problem was proposed by Hochreiter et al. [16] by using a variation of RNNs called Long-Short Term Memory (LSTM). LSTMs use a memory cell (*c_t_*), to keep track of long-term relationships between text. Using a gated architecture (input, output, and forget), LSTMs are able to modulate the exposure of a memory cell by regulating the gates. LSTMs can be defined as:

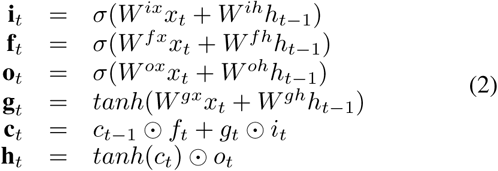

where, **i***_t_*, **f***_t_*, and **o***_t_* are the input, forget, and output gates respectively. Each gate uses a *sigmoid* (*σ*) function applied over the sum of input *x_t_* and previous hidden state *h_t_*_−1_ (multiplied with their weight matrices *W*). **g***_t_* denotes the candidate state computed over a *tanh* function on the input and previous hidden state. *W^ix^*, *W^fx^ W^ox^ W^gx^* are weight matrices used with input *x_t_*, while *W^ih^*, *W^fh^*, *W^oh^*, and *W^gh^* are used with hidden states for each gate and candidate state. The memory cell **c***_t_* utilizes the forget gate (**f***_t_*) and multiplies (⊙ - element-wise) it old memory cell **c***_t_*_−1_ and adds to the state of candidate (**g***_t_*) multiplied with the input gate (**i***_t_*). The hidden state is given by a *tanh* function applied to the memory cell *c_t_* multiplied with output gate (**o***_t_*).

#### 4) Gated Recurrent Unit

A variation on LSTM, was introduced by Cho et al. [17] as Gated Recurrent Unit (GRU). Using update and reset gates, GRUs are able to control amount of information within a unit (without a separate memory cell as with LSTM). GRUs can formally be defined as

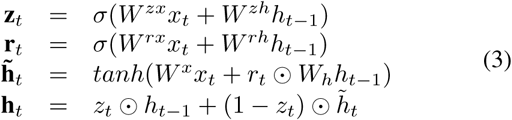

where, **z***_t_* and **r***_t_* are update and reset gates respectively, 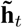 is the candidate activation/hidden state.

Similar to the LSTM architecture, GRUs benefit from the additive properties in their network to remember long term dependencies, and solve the vanishing gradient problem. Since GRUs do not utilize an an output gate, they are able to write the entire contents of their memory cell to the network. The lack of a memory cell also makes GRUs more efficient in comparison to LSTMs.

### D. Deep learning Architectures

Below, we describe three architectures - window-based, word-based, and word+character based to be used in conjunction with the different models described above.

#### 1) Window-based

In this architecture, the window-based input (*i_v_*) consists of feature vectors (*f_v_*) for each word/term (*t*) within an encoded sentence. Each *f_v_* consisted of the following attributes:

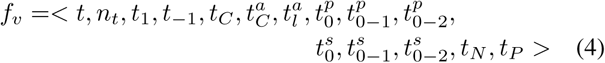

where,

*t* is the term,

*n_t_* is the number of terms in the sentence,

*t*_1_ is a boolean value indicating if the term is the first term in the sentence,

*t*_−1_ is 1 if term is the last term in the sentence 0 otherwise,

*t_C_* is 1 if first letter in *t* is uppercase,

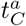 is 1 if all letter in *t* are uppercase,

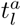 is 1 if all in *t* are lower case,

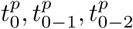 record character prefixes of *t* at various window size,

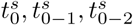 record character suffixes of *t* at various window sizes,

*t_N_* and *t_P_* are the next and previous terms respectively.

#### 2) Word-based

Each word and its corresponding annotation labels (tags) are encoded with integer values, derived from unique words and annotations present in the corpus. The dataset was based on unigram annotations that only use ontology annotations where a single word in text maps to an ontology concept.

In word-based architectures (Figure 1), the input 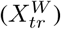 is provided to an Embedding layer which converts the input into dense vectors of 100 dimensions. The output vectors are then fed to a bidirectional model (RNN/GRU/LSTM) consisting of 150 hidden units. The output from the model goes to a dense perceptron layer using ReLU activation which also employs a 0.6 Dropout. The output is further fed into a CRF layer which looks for correlations between annotations in close sequences to generate the predictions (*y_pr_*).

**Fig. 1.**
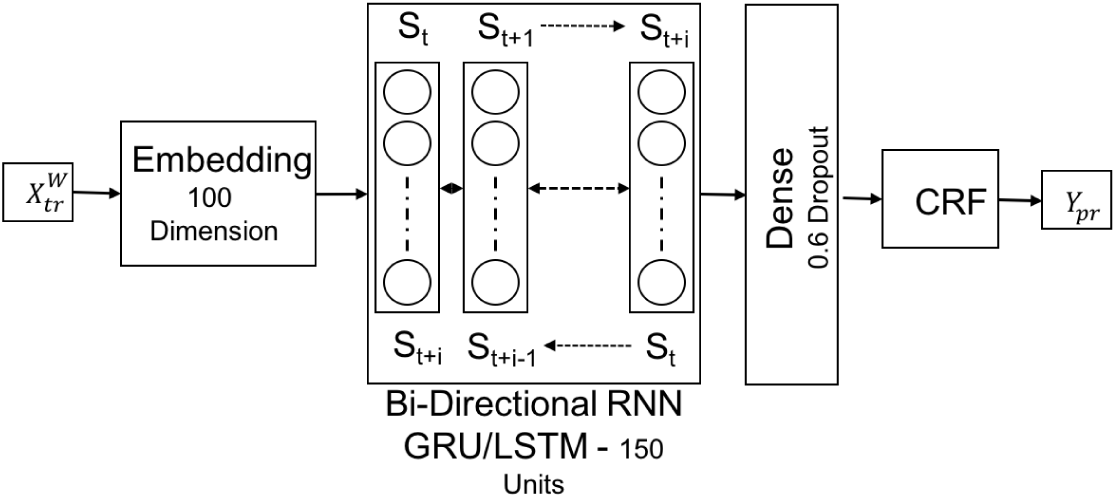
Word-based architecture using bidirectional RNN/GRU/LSTM models

#### 3) Character+Word Based

A Character+Word based architecture is similar to the word based architecture described above. In addition to word-based inputs 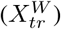 is also takes advantage of characters 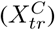 within words to make predictions.

### E. Model development

We developed a new deep learning architecture that uses a Character+word based architecture coupled with two bidirectional Gated Recurrent Units. Our architecture (Figure 2) consists of character level input 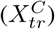 provided to an Embedding layer (*E*_1_) which compresses the dimensions of characters to the number of unique annotations in the corpus (*NT ags*).

**Fig. 2.**
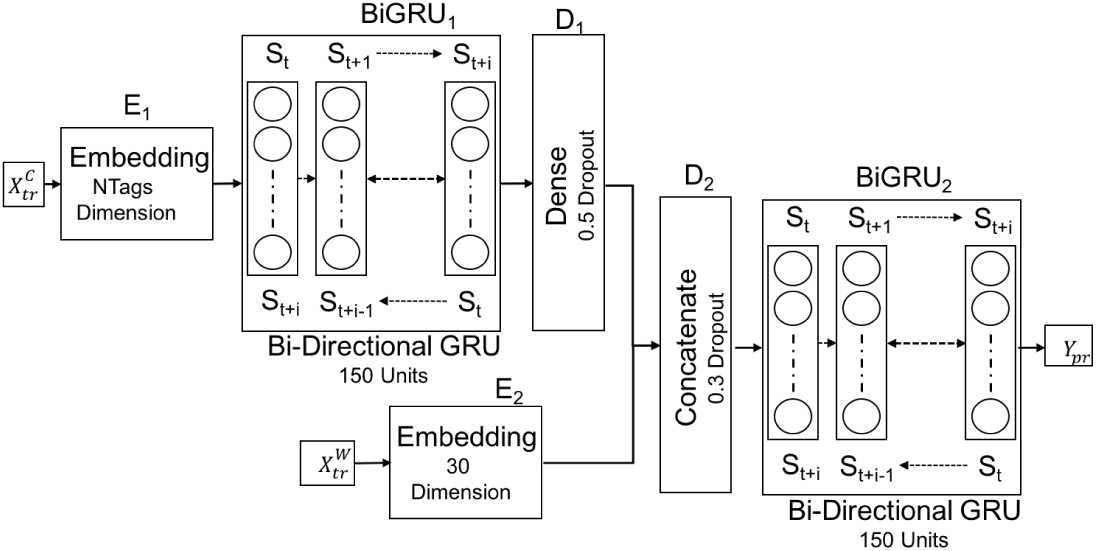
Character+word based architecture using two bidirectional GRU models.

The output of Embedding layer *E*_1_ is fed to a bidirectional GRU (*BiGRU*_1_) layer with 150 units followed by a 60% output drop in a Dropout layer (*D*_1_). Simultaneously, the word-level input 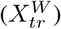 was provided to a second Embedding layer (*E*_2_) with 30 dimensions. The output from *E*_2_ was concatenated with the output from the first Dropout layer *D*_1_ and fed through a second Dropout layer (*D*_2_) with a 30% drop. Output from *D*_2_ was fed into a second bidirectional GRU layer (*BiGRU*_2_) consisting of 150 units.

The above model was tested with and without a final *CRF* layer leading to two new configurations - *CW* − *BiGRU* − *CRF* and *CW* − *BiGRU*. The models were run for 15 epochs with a batchsize of 32 instances.

### F. Model Comparison

We compared the performance of our new Character+word based GRU architecture and the two models developed therein (*CW* − *BiGRU* − *CRF*, *CW* − *BiGRU*) (Section IV-E) to six state of the art models that have been used in prior work. Below, we specify the component details of each of the six prior models that have been evaluated.

#### 1) MLP

Multi layer perceptrons were used with a window based architecture to create a three layered (input, hidden, output) *MLP* model. The input and the hidden layer consisted of 512 perceptrons with a Rectified Linear Unit (ReLU) activation function while the output layer consisted of perceptrons equal to the number of unique annotations in the corpus (*NT ags*). 20% Dropout was used for the hidden and output layers to prevent overfitting of the data. Categorical cross-entropy was used for calculating the loss function and NAdam (Adam RMSprop with Nesterov momentum) was used as the optimizer function. Each of the feature vectors (from the training data), were fed into the MLP architecture for 15 epochs with a batch size of 256.

#### 2) BiRNN-CRF

The BiRNN-CRF model uses a word-based input coupled with a BiRNN model and ending with a CRF model. Similar to the BiRNN architecture (Figure 1), the BiRNN-CRF model consists of a 100 dimension Embedding layer followed by a BiRNN with 150 units followed by a 0.6 Dropout layer. The output of the 0.6 Dropout layer is fed to a CRF which generated the predicted output.

#### 3) BiLSTM-CRF

The BiLSTM-CRF model is identical to the BiRNN-CRF except that it uses a LSTM in place of the RNN.

#### 4) BiGRU-CRF

The BiGRU-CRF model is identical to BiRNN-CRF and BiLSTM-CRF except that it uses a Gated Recurrent Unit in place of the RNN or LSTM.

#### 5) CW-BiLSTM

The CW-BiLSTM model is similar to the CW-BiGRU model described above (see Section IV-E) except that the BiGRU is replaced with a BiLSTM.

#### 6) CW-BiLSTM-CRF

The CW-BiLSTM-CRF model is developed by adding a CRF layer at the end of the CW-BiLSTM model pipeline indicating that the output of the CW-BiLSTM model would be fed to a CRF layer to generate the final predictions.

### G. Parameter Tuning

The GO annotation data was split into training and test sets using a 70:30 ratio. The training set was used to tune the following parameters for all models. Multiple architecture parameters such as - 1) Number of layers in MLP (along with number of perceptrons), 2) Number of units in RNN/GRU/LSTM, 3) Embedding Dimensions for Characters and Words, and 4) Optimization functions, were evaluated for model performance. A grid-search model was explored, where each architecture was evaluated for different combinations of the parameter. In each case, model performance metrics were recorded in form of Precision, Recall, F1-score, and Jaccard similarity.

### H. Experiments to predict ontology annotations

The largest number of annotations in the CRAFT corpus came from the Gene Ontology. So, we first used the GO annotations to train and test the suite of 8 models described above. Subsequently, we applied the best model from these experiments to annotate the CRAFT corpus with the other four ontologies (Chebi, Cell, Protein, and Sequence corpora).

Root-Mean-Square propagation (RMSProp) optimizer was used to test the performance of the different models. A batch size of 32 along with 15 epochs was used for model training. Performance characteristics in terms of train-test loss (calculated using the CRF function), prediction precision, recall, F1-score along with mean semantic similarity score was recorded for each model.

## V. Results and Discussion

The CRAFT corpus contains 67 full length papers with annotations from five ontologies (GO, CHEBI, Cell, Protein, and Sequence). For each of these ontologies, we extracted all sentences across the 67 papers with at least one annotation for the ontology. The largest number of annotations came from the GO (Table I) while the Cell ontology accounted for the lowest number of annotations.

**TABLE I.** Characteristics of the Craft corpus - Number of sentences with at least one annotation, number of unique annotations (unigrams only), and number of unique words in the corpus.

| Dataset | Number of Sentences | Number of Unique Annotations | Number of Unique Words in the Corpus |
| --- | --- | --- | --- |
| GO | 17,921 | 359 | 9,571 |
| Sequence | 15,606 | 156 | 7,262 |
| Protein | 12,621 | 546 | 5,153 |
| Chebi | 11,109 | 309 | 3,127 |
| Cell | 9,088 | 68 | 3,042 |

Figure 3 shows the loss and accuracy trends for each model on the GO annotation data. The goal of the models is to minimize loss while increasing accuracy as the number of epochs increase.

**Fig. 3.**
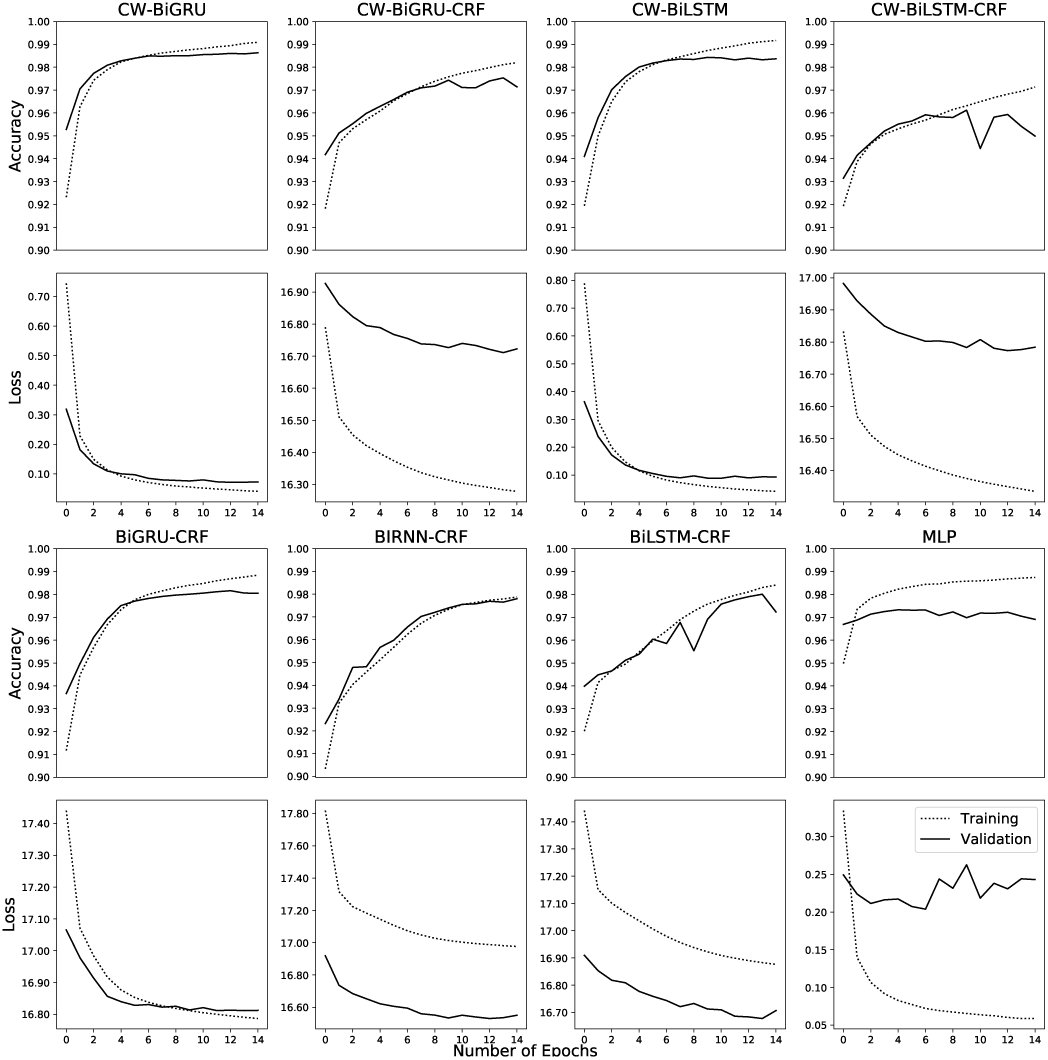
Comparison of model loss and accuracy on training and validation data using Gene Ontology annotations

First, we see that our CW-BiGRU model shows improvement in both training and validation accuracy as the number of epochs increase. Correspondingly, we observe a decrease in training and validation loss indicating that the model is able to self-improve with each subsequent epoch.

The CW-BiGRU-CRF model initially shows the same accuracy improvement like the CW-BiGRU model but later increases in epochs result in a divergence in the training and validation accuracy indicating that the model might be prone to overfitting. While there is a substantial decrease in training loss, a similar decrease is not observed in validation loss.

CW-BiLSTM shows similar trends to CW-BiGRU. CW-BiLSTM-CRF training and validation accuracy increase similarly until a certain point after which the validation accuracy drops and diverges sharply from the training curve indicating a case of overfitting.

BiGRU-CRF and BiRNN-CRF models show substantial improvement in accuracy with increasing epochs. However, BiRNN-CRF shows divergence in the loss patterns. Similar to CW-BiLSTM-CRF, BiLSTM-CRF also shows signs of over-fitting in the accuracy patterns. MLP is the worst performing model with very minor improvements in validation accuracy as the number of epochs increase indicating that the model is unable to improve itself with each subsequent epoch.

It is clear that the CW-BiGRU models are able to outperform the other models by improving accuracy and reducing loss with each epoch without overfitting.

A large proportion of input data is not annotated to GO terms but to a tag ‘*O*’ indicating the absence of an annotation. In addition to accurately predicting GO annotations, the model also needs to accurately predict the absence of an annotation. However, given the disproportionate amount of data pertaining to the absence of annotations, the models were observed to predict the absence of annotations remarkably accurately in comparison to predicting presence.

To provide a more conservative view of the models’ performance, we report Precision, Recall, F-1 Score, and Jaccard similarity (Table II) only on data indicating presence of ontology terms, i.e. text annotated with an ontology term. Unlike the accuracy measurements above, the metrics below do not take into account the models’ performance at identifying the absence of annotations, but rather focus on ability to identify annotations when they’re present in the Gold Standard.

**TABLE II.**
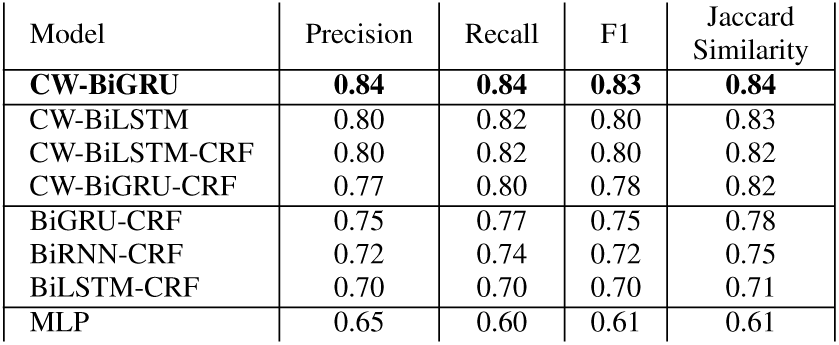
Precision, Recall, F1, and Jaccard Similarity scores for the eight models on CRAFT Gene Ontology annotation data.

These results (Table II and Figure 3) show that our model (CW-BiGRU) outperforms the other 7 models in all four metrics. Our model outperforms the best among the other 7 models (CW-BiLSTM) by 4% (Precision), 2% (Recall), 3% (F1 score), 1% (Jaccard similarity).

Additionally, we observe that character-word based models (CW-BiGRU, CW-BiLSTM, CW-BiLSTM-CRF, CW-BiGRU-CRF,) outperform models that use only word embeddings.

Among the character-word based models, surprisingly, the addition of an extra CRF layer (CW-BiLSTM-CRF, CW-BiGRU-CRF) either fails to improve performance (e.g CW-BiLSTM vs. CW-BiLSTM-CRF) or leads to a decline in performance (e.g CW-BiGRU vs. CW-BiGRU-CRF) as compared to not using a CRF end layer (CW-BiLSTM, CW-BiGRU). The MLP model shows substantially lower performance as compared to the other models across all four metrics. The Accuracy and Loss plots (Figure 3) suggest that the decline in performance when adding a CRF layer is due to potential overfitting.

We explored how predictions from our best model, CW-BiGRU, diverge from the Gold Standard. We found that the majority of predictions (89.25%) are an exact match for the CRAFT annotations. Surprisingly, only a small proportion of predictions are partial matches (2.45%). 8.26% of the model’s predictions are false negatives while 6.38% are false positives. We hypothesize that one of the primary reasons for false negatives might be lack of enough training instances for those particular GO annotations.

Finally, we applied the best performing model from the above evaluation (CW-BiGRU) and tested it on data from four other ontologies. Interestingly, the model shows better prediction performance on the other ontologies as compared to GO despite the substantially smaller training datasets (Table III).

**TABLE III.** Precision, Recall, F1, and Jaccard Similarity scores for the eight models on annotations from five ontologies in CRAFT.

| Model | Ontology | Precision | Recall | F1 | Jaccard Similarity |
| --- | --- | --- | --- | --- | --- |
| CW-BiGRU | Cell | 0.92 | 0.92 | 0.92 | 0.925 |
| CW-BiGRU | Protein | 0.91 | 0.90 | 0.90 | 0.917 |
| CW-BiGRU | CHEBI | 0.86 | 0.87 | 0.86 | 0.882 |
| CW-BiGRU | GO | 0.84 | 0.84 | 0.83 | 0.843 |
| CW-BiGRU | Sequence | 0.83 | 0.86 | 0.84 | 0.864 |

## VI. Conclusions and Future Work

The data used in this study was limited to single words annotated to ontology concepts (unigrams). Next, we will explore more robust models including n-grams to account for sequences of words tagged with an annotation. Future work will also include models that can be trained to weight the prediction of some target classes higher than others. These models would be able to prioritize presence prediction of annotations as compared to the absence of an annotation.

This study demonstrates the utility of deep learning approaches for automated ontology-based curation of scientific literature. Specifically, we show that models based on Gated Recurrent Units are more powerful and accurate at annotation prediction as compared to the LSTM based models in prior work. Our findings indicate that deep learning is a promising new direction for ontology-based text mining, and can be used for more sophisticated annotation tasks (such as phenotype curation) that build upon Named Entity Recognition.

